# The physiological impact of an N-terminal Halo-tag on GIPR function in mice

**DOI:** 10.1101/2024.12.18.629103

**Authors:** Iona Davies, Yusman Manchanda, Kyle W Sloop, Stephen R Bloom, Tricia MM Tan, Alejandra Tomas, Ben Jones

## Abstract

**Aims:** G protein-coupled receptors (GPCRs) modified with self-labelling enzymatic tags (e.g. Halo, SNAP, CLIP) have enabled the *in vitro* study of receptor expression and trafficking. We developed a *Gipr^Halo/Halo^* mouse model to gain insights into endogenous glucose-dependent insulinotropic polypeptide (GIP) receptor (GIPR) signalling, an important modulator of glucose and appetite homeostasis.

**Methods:** *Gipr^Halo/Halo^* mice were generated through ES cell-based gene targeting and physiologically characterised via body weight, oral glucose tolerance test and intraperitoneal glucose tolerance test analysis. Live cell imaging was used to measure cAMP responses in dispersed pancreatic islets from *Gipr^Wt/Wt^* and *Gipr^Halo/Halo^* littermates in response to GIP. cAMP accumulation and surface receptor expression were assessed in AD293 cells transiently transfected with mouse and human untagged GIPR and Halo-GIPR. Endogenous pancreatic islet expression of Halo-GIPR was determined by immunohistochemistry.

**Results:** *Gipr^Halo/Halo^* mice displayed comparable body weights and responses to oral glucose administration as *Gipr^Halo/Wt^* and *Gipr^Wt/Wt^* littermates. However, *Gipr^Halo/Halo^* mice only responded to high dose human GIP during intraperitoneal glucose tolerance testing, and cAMP responses to GIP were impaired in *Gipr^Halo/Halo^* islets. Despite unimpaired surface expression, *in vitro* cAMP responses were lessened in AD293 cells expressing Halo-GIPR compared to untagged GIPR. Anti-Halo staining in mouse *Gipr^Halo/Wt^* pancreatic islets displayed detectable signal.

**Conclusions:** While the *Gipr^Halo/Halo^* mouse model could be used to study endogenous GIPR expression, due to reduced responses we advise against its use for endogenous GIPR trafficking and signalling. Our study highlights the importance of carefully phenotyping mouse models in which modifications to endogenous proteins have been introduced.

## Introduction

The role of glucose dependent insulinotropic polypeptide (GIP) signalling in energy and glucose homeostasis is a hot area of metabolic research. The study of GIP receptor (GIPR) expression and trafficking, vital aspects in our understanding of GIPR physiology and pharmacology, have been hindered by the lack of commercially available validated GIPR antibodies, meaning that many studies have relied on the presence of *Gipr* mRNA to determine GIPR expression in both mouse and human tissue (1–4). The development of fluorophore-conjugated GIPR agonists has provided some insights into endogenous GIPR distribution and behaviour (4,5), but provide only indirect evidence of receptor localisation and may be misleading in some contexts, e.g. if agonists dissociate from receptors after entering the endocytic pathway.

G protein-coupled receptors (GPCRs) modified with self-labelling enzymatic tags such as SNAP, Halo and CLIP are becoming increasingly common in pharmacological research as a method of studying receptor expression and trafficking *in vitro* (6). These systems rely on the establishment of sensitive and specific interactions between these enzyme tags and their corresponding substrate probes, detectable via their conjugation to fluorescent dyes or other types of chemical labels. The SNAP-tag was the first enzyme self-label to be developed (7,8). This is an evolved version of the DNA repair protein O6-alkylguanine DNA alkyltransferase which reacts with O6-benzylguanine to form an irreversible covalent bond and liberate guanine. The Halo-tag – a modified bacterial haloalkane dehalogenase which reacts with chloroalkanes to form a covalent alkyl-Halo-tag product and liberate chloride ions - was developed shortly afterwards (9). The use of SNAP- or Halo-tagged GIPR and glucagon-like peptide-1 receptor (GLP-1R) constructs is now commonplace in the study of *in vitro* incretin receptor trafficking (10,11).

Advances in genetic engineering have enabled the generation of mice which endogenously express proteins tagged with enzyme self-labels, enabling protein expression and trafficking to be investigated *in vivo*. In 2023, a characterisation of the *Glp1r^SNAP/SNAP^* mouse model was published (12). Investigators performed sophisticated experiments studying pancreatic islet GLP-1R trafficking and found that constitutive internalisation was higher than reported using cell lines, emphasising the importance of studying receptor behaviours in their native environment.

To this end, we developed a *Gipr^Halo/Halo^* mouse model for the study of endogenous GIPR expression and trafficking. Our aim was to explore central and pancreatic GIPR expression and trafficking in response to chronic GIPR agonism, relevant in deciphering why both GIPR agonists and antagonists reduce body weight, as well as whether the GIPR is a suitable candidate for the development of agonists biased away from GIPR internalisation. The Halo-tag was selected as it could provide the flexibility to study GIPR alongside other SNAP-tagged GPCRs in the same animal in future projects. However, physiological analysis of this mouse model showed that Halo-tagging the GIPR partially impaired receptor function, thus limiting its uses beyond the study of GIPR expression. We therefore conclude that caution should be applied when using or developing mouse models with enzyme-self labelled endogenous proteins, as these methods might result in unforeseen negative consequences impacting normal protein function.

## Methods

### Peptides

Mouse GIP(1–42) and human GIP(1–42) were purchased from Bachem, Switzerland. GIP-TMR, a previously described fluorescent analogue of human GIP(1–42) (13), was purchased from Wuxi AppTec, China.

### Cell culture

AD293 cells (Agilent, US) were maintained in Dulbecco’s modified medium (DMEM, Thermo Fisher, UK) with 1% penicillin/streptomycin (P/S, Sigma, UK) and 10% foetal bovine serum (FBS, Thermo Fisher, UK) (complete DMEM).

### Plasmids

Plasmids encoding full length wild-type human and mouse GIPR in the pcDNA/FRT vector (Thermo Fisher, UK), and human and mouse GIPR featuring an N-terminal extracellular Halo-tag in the pcDNA5/FRT vector were custom synthesised by Genewiz, UK.

### Homogenous Time Resolved Fluorescence (HTRF) cAMP accumulation assay

HTRF cAMP accumulation assays were performed using the cAMP-Gs Dynamic 2 kit (Cisbio, France) as per the manufacturer’s protocol in AD-293 cells 24 hours after transient transfection with specified plasmid. Full details have been described previously (14).

### Cell surface labelling

24 hours post transient transfection with specified plasmid, AD-293 cells were labelled for 1 hour with GIP-TMR (100 nM) in DMEM + 0.1% bovine serum albumin (BSA), prior to fixation with 1% paraformaldehyde and imaging using an epifluorescence microscope.

### *In vivo* studies – husbandry

All animal procedures were approved by the British Home Office UK Animals (Scientific Procedures) Act 1986. All experiments were performed using both female and male *Gipr^Wt/Wt^*, *Gipr^Halo/Wt^* and *Gipr^Halo/Halo^* 6–24-week-old littermates. For all *in vivo* studies, mice were group housed in individually ventilated cages with a standard 12:12 hour light-dark cycle. Unless fasted, mice had free access to food (RM1-Special Diet Services, UK) and water.

### Generation of *Gipr^Halo/Halo^* mice

*Gipr^Halo/Halo^* mice generation was outsourced to Cyagen. ES cell-based gene targeting was employed. The Halo tag cassette was inserted into the *Gipr* allele after the signal peptide in exon 2 and *loxP* sites were inserted either side of exon 5. A neomycin resistance cassette was chosen as the positive selection marker and diptheria toxin A as a negative selection marker. The targeting construct was electroporated into C57BL/6N ES cells and southern blot analysis was carried out to identify PCR-positive clones. Clones were then micro-injected into host embryos and transferred into surrogate mothers. Following subsequent chimera breeding and identification of heterozygous pups, sperm samples from two 11-week-old male heterozygous mice were cryopreserved and subsequently transferred to ES Cell & Transgenic Facility of the Laboratory of Medical Sciences, Medical Research Council where the mouse line was rederived using *in vitro* fertilisation.

### Genotyping

Genotyping was performed using the KAPA Mouse Genotyping Kit (Sigma, UK) as per the manufacturer’s instructions. To identify the presence of the Halo-tag, a primer set for the Halo construct (forward: GATCCCAGCCTCACTTATCTACTG, reverse: GACCAACATCGACGTAGTGCAT) (expected fragment of 364 base pairs) and a primer set for a constitutively expressed cDNA sequence (forward: GCAGAAGAGGACAGATACATTCAT, reverse: CCTACTGAAGAATCTATCCCACAG) (hexokinase, expected fragment of 689 base pairs) were used. To identify the presence of the *loxP* sites between exon 5 (and thus confirm whether a mouse is *Gipr^Halo/Wt^* or *Gipr^Halo/Halo^*), a primer set was used where the wild-type exon 5 allele has an expected fragment of 281 base pairs and the *loxP*-flanked exon 5 allele has an expected fragment of 337 base pairs (forward: TGGGCTCTGTCACATGATTTACTTA, reverse: GAGGGTGTCAGAGGTGTAGC).

### Intraperitoneal glucose tolerance tests (IPGTTs)

Mice were fasted at 08:00 hours prior to intraperitoneal glucose injection ± human GIP at a specified dose at 14:00 hours. The injection volume was adjusted to the body weight of the mouse so that all mice received glucose at 2 g/kg. Blood glucose measurements were taken via tail venesection at t=0, t=20, t=40 and t=60 minutes. Glucose readings were measured in mmol/L using a GlucoRx glucometer.

Three litters of mice were used for these studies: cohort 1 (*Gipr^Wt/Wt^*, male n=10, female n=10; *Gipr^Halo/Wt^*, male n=13, female n=8; *Gipr^Halo/Halo^*, male n=4, female n=4), cohort 2 (*Gipr^Wt/Wt^*, male n=1, female n=3; *Gipr^Halo/Wt^*, male n=4, female n=8; *Gipr^Halo/Halo^*, male n=1, female n=3) and cohort 3 (*Gipr^Wt/Wt^*, male n=2, female n=1; *Gipr^Halo/Wt^*, male n=1, female n=2; *Gipr^Halo/Halo^*, male n=4, female n=2).

For studies testing GIP (50 nmol/kg), cohorts 1 and 2 were used, giving the following totals for each genotype; *Gipr^Wt/Wt^* male n=11, female n=13; *Gipr^Halo/Wt^*, male n=17, female n=16; *Gipr^Halo/Halo^*, male n=5, female n=7.

For studies testing GIP (200 nmol/kg), some of cohort 1 and all of cohort 2 and 3 were used, giving the following totals for each genotype; *Gipr^Wt/Wt^*, male n=5, female n=6; *Gipr^Halo/Wt^* male n=7, female n=13; *Gipr^Halo/Halo^* male n=7, female n=7.

For studies testing GIP (500 nmol/kg), some of cohort 1 and all of cohort 2 and 3 were used, giving the following totals for each genotype; *Gipr^Wt/Wt^* male n=5, female n=6; *Gipr^Halo/Wt^*, male n=7, female n=13; *Gipr^Halo/Halo^* male n=7, female n=7.

Of note, the study conducted with cohort 3 was a three-way crossover study administering vehicle, GIP (200 nmol/kg) and GIP (500 nmol/kg) with glucose injection. Thus, when data was combined to separately compare vehicle *versus* GIP (200 nmol/kg) and vehicle *versus* GIP (500 nmol/kg) across all cohorts, the data collected from the saline group of cohort 3 was used in both analyses.

### Oral glucose tolerance tests (OGTTs)

Cohort 1 was used for OGTTs. Mice were fasted at 08:00 hours prior to oral gavage of glucose at 14:00 hours, with the injection volume adjusted to the body weight of the mouse so that all mice received glucose at 2 g/kg. Blood glucose measurements were taken via tail venesection at t=0, t=20, t=40 and t=60 minutes. Glucose readings were measured in mmol/L using a GlucoRx glucometer.

### *Ex vivo* islet dose responses

Pancreatic islets from *Gipr^Wt/Wt^* and *Gipr^Halo/Halo^* mice were isolated as previously described (14), dispersed into single cells via 3-minute trituration with warm 0.05% trypsin-EDTA, and resuspended in RPMI-1640 (Thermo Fisher, UK) + 10% FBS + 0.1% P/S. Cells were transduced with the Green Up cADDis biosensor (Molecular Montana, US) (15) prior to seeding on 96-well plates coated with 0.01% poly-D-lysine hydrobromide and 25 µg/mL mouse laminin. Following overnight incubation (37°C, 95%:5% O_2_ :CO_2_ ratio), cells were imaged in Krebs-Ringer Bicarbonate buffer (140 mM NaCl, 3.6 mM KCl, 1.5 mM CaCl_2_, 0.5 mM MgSO_4_, 0.5 mM NaH_2_PO_4_, 2 mM NaHCO_3_, 10 mM HEPES, saturated with 95% O_2_/5% CO_2_, pH 7.4) with 6 mM glucose and 0.1% BSA at 37°C using an automated epifluorescence microscope, enabling multiple fields of view to be captured in parallel. Acquisitions were 36 minutes in length, consisting of a 3-minute baseline recording pre-peptide addition, 6 x 5- minute recordings following addition of 10-fold increasing concentrations of peptide and a final 3- minute recording following addition of IBMX (500 µM) and forskolin (50 µM) to maximally stimulate the sensor. cADDis signal quantification was performed using Fiji v1.54f (NIH), where islet cell fluorescence was normalised to both baseline and IBMX/FSK responses, and the area under the curve (AUC) calculated for each cell.

### *Ex vivo* islet immunohistochemistry

Pancreatic islets from *Gipr^Wt/Wt^*,*Gipr^Halo/Wt^* and *Gipr^Halo/Halo^* mice were isolated. Islets were fixed in paraformaldehyde (4%) and blocked without permeabilization in 1% BSA prior to 48-hour incubation with anti-Halo antibody (1:500; Promega, UK) in 0.1% BSA/PBS at 4°C. Islets were then washed with 3xPBS and incubated with secondary anti-rabbit AlexaFluor 568 (1:500; Thermo Scientific, UK). Islets were imaged using a Leica Stellaris 8 inverted confocal microscope with a 63x/1.40 oil objective with Lightning super-resolution modality from the Facility for Imaging by Light Microscopy (FILM) at Imperial College London.

### Statistical analysis

Analyses was conducted using Prism 10.0 (GraphPad software). Statistical significance was calculated using one sample t-test, one- or two-way ANOVA, as indicated in the figure legends. Šídák and Dunnett’s tests were used to correct for multiple comparisons. Unless specified otherwise, all summarised data points are presented as mean□±□SEM. For concentration response experiments, 3-parameter fits are plotted. Statistical significance was determined as *P<0.05, **P<0.01, ***P<0.001 and ****P<0.0001.□□

## Results

### Generation of the *Gipr^Halo/Halo^* mouse model

N-terminal tags facilitate cell surface-specific labelling (**Figure 1A**), and previous studies using heterologous expression systems show that GIPR tolerates N-terminal Halo- and SNAP-tag modifications (13,16), as long as the tag is not upstream of the endogenous GIPR signal peptide. To adapt this approach to natively expressed GIPR, we designed a *Gipr^Halo/Halo^* mouse model, in which the sequence for the Halo-tag protein was inserted into the *Gipr* allele downstream of the Gipr signal peptide in exon 2, and *loxP* sites inserted between exon 5 (**Figure 1B**) via ES-cell based gene targeting (**Figure 1C**). *Gipr^Halo/Wt^* breeding lines were established to produce multiple litters of *Gipr^Wt/Wt^*, *Gipr^Halo/Wt^* and *Gipr^Halo/Halo^* mice (**Figure 1D**).

**Figure 1:**
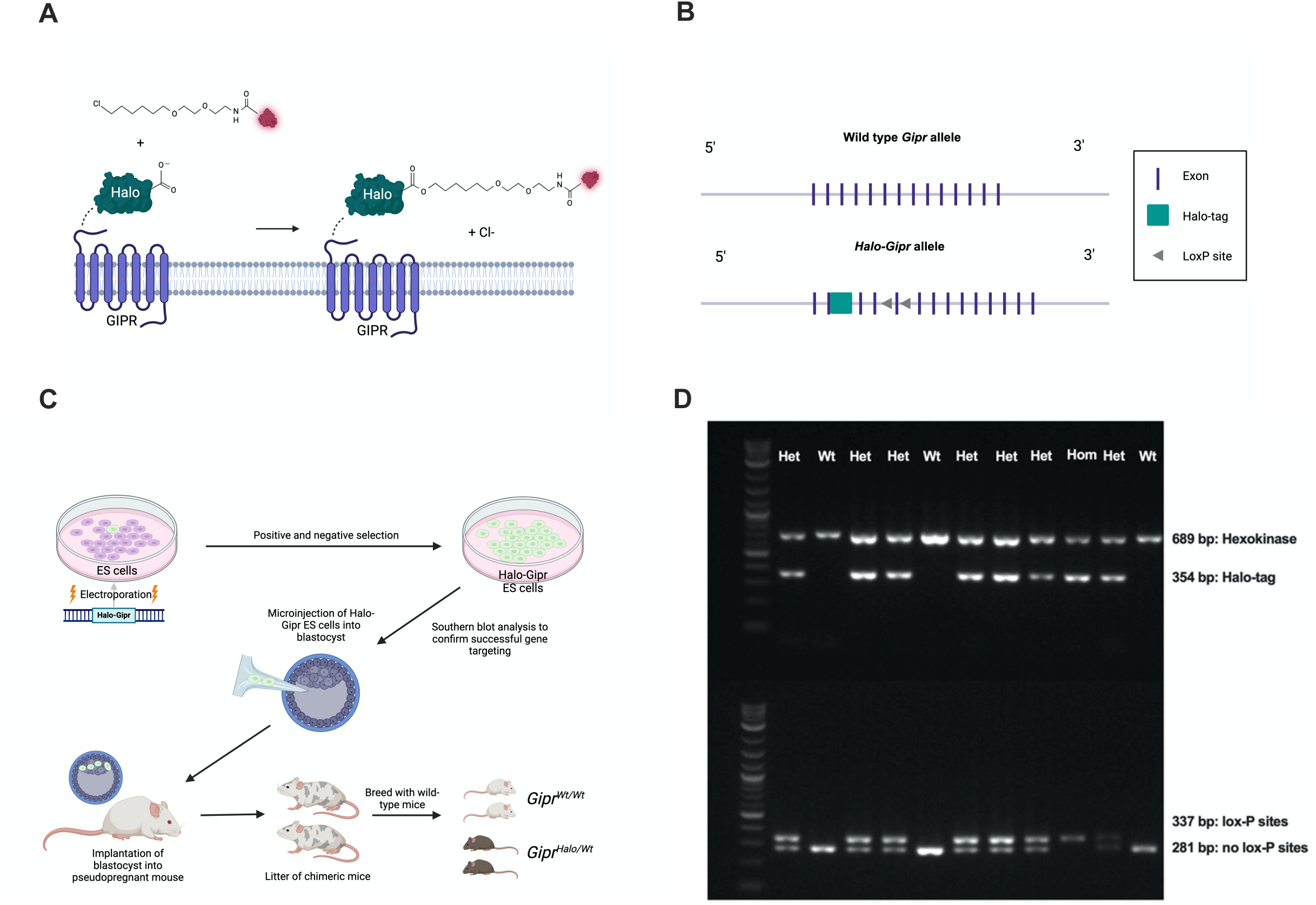
Generation of the *Gipr^Halo/Halo^* mouse model. **(A)** Schematic of GIPR tagging with Halo-tag protein and subsequent receptor labelling. (**B**) Wild-type *Gipr* and *Halo-Gipr* alleles. (**C**) ES cell-based gene targeting used to generate the *Gipr^Halo/Halo^* mouse model. (**D**) Example agarose gels showing presence of hexokinase DNA (689 bp) and Halo-tag DNA (354 bp) (top), and lox-P sites at *Gipr* exon 5 (337 bp) or wild-type exon 5 (281 bp) (bottom). 1 kb Plus DNA ladder shown at the left-hand side. The genotype derived from band presence is displayed at the top of the figure. Wt = *Gipr^Wt/Wt^*, Het = *Gipr^Halo/Wt^*, Hom = *Gipr^Halo/Halo^*.

### *Gipr^Halo/Halo^* mice show impaired anti-hyperglycaemic responses to GIP

To establish whether Halo-tagging affects physiological GIPR signalling, we investigated differences in body weight gain and handling of oral glucose administration between Halo-tagged and wild-type genotypes. There were no statistically significant differences in body weight between genotypes of either sex at 6 weeks (**Figure 2A, B**), or between 6-9 weeks, measured weekly in a subset of mice (**Figure S1**). However, this does not necessarily preclude central GIPR signalling disturbances as body weight differences between *Gipr* KO and wild-type mice are only revealed upon high fat diet feeding (17,18). Plasma glucose responses to an oral glucose bolus were identical across all genotypes, suggesting that physiological incretin capacity was preserved in *Gipr^Halo/Halo^* and *Gipr^Halo/Wt^* mice (**Figure 2C, D**).

**Figure 2:**
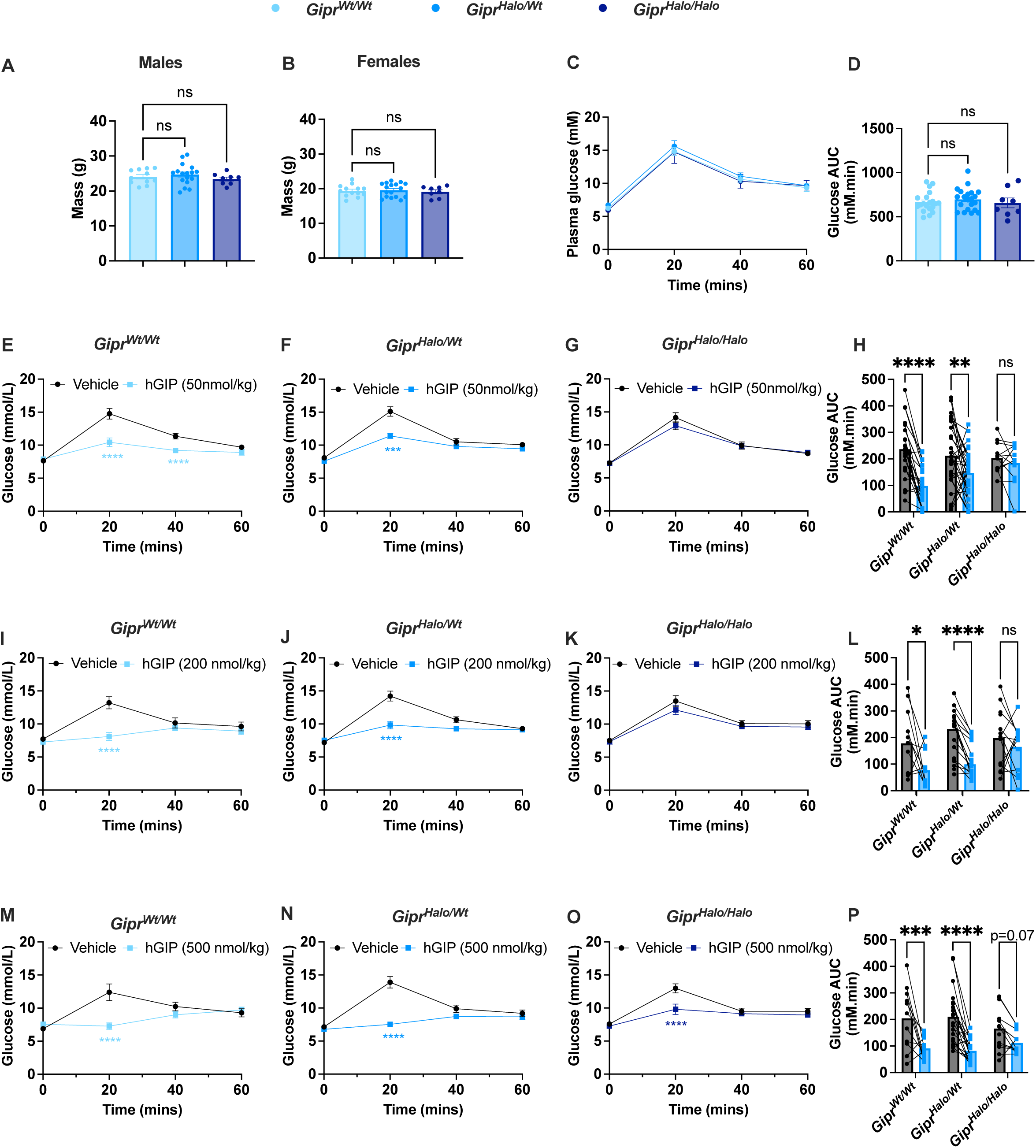
*Gipr^Halo/Halo^* mice improve glucose tolerance in response to higher doses of human GIP than Gipr*^Wt/Wt^* mice. **A**, **B**) Body mass (g) of *Gipr^Wt/Wt^* (male: n=11, female: n=13), *Gipr^Halo/Wt^* (male: n=17, female: n=17) and *Gipr^Halo/Halo^*(male: n=8, female: n=8), measured at 6 weeks of age. Males are displayed in **A** and females in **B**. **C**, **D**) OGTT conducted in *Gipr^Wt/Wt^* (n=20), *Gipr^Halo/Wt^* (n=21) and *Gipr^Halo/Halo^* (n=8) mice. **E**-**P**) Crossover IPGTTs conducted in *Gipr^Wt/Wt^* mice (**E**: n=24, **I**: n=11, **M**: n=11), *Gipr^Halo/Wt^* mice (**F**: n=33, **J**: n=20, **N**: n=20) and *Gipr^Halo/Halo^* mice (**G**: n=12, **K**: n=14, **O**: n=14), in response to human GIP (hGIP, 50 nmol/kg) (**E**-**G**), hGIP (200 nmol/kg) (**I**-**K**) and hGIP 500 nmol/kg (**M**-**O**). **C**, **E**-**G**, **I**-**K**, **M**-**O**) Plasma glucose time-course. **D**, **H**, **L**, **P**) Glucose AUC derived from corresponding glucose curves. Blood glucose at specific time-points were analysed using a two-way ANOVA with time and subgroup as co-variables. The Šídák test used to correct for multiple comparisons. Body mass at 6 weeks (**A**, **B**) and glucose AUC in **C** were analysed with a one-way ANOVA. The Dunnett’s test was used to correct for multiple comparisons. Glucose AUCs in **H**, **L** and **P** were analysed using a two-way ANOVA with genotype and subgroup as co-variables. The Šídák test was used to correct for multiple comparisons. All values are presented as a mean ± SEM. *P<0.05, **P<0.01, ***P< 0.001, ****P< 0.0001.

To probe GIP responses more specifically, we tested responses to increasing doses of human GIP during an IPGTT. Here, while human GIP (50 nmol/kg and 200 nmol/kg) significantly improved glucose tolerance in *Gipr^Wt/Wt^* and *Gipr^Halo/Wt^* mice, neither dose was able to improve glucose tolerance in *Gipr^Halo/Halo^* mice, suggestive of impaired pancreatic GIPR functionality (**Figure 2E-L**). *Gipr^Halo/Halo^* mice did respond to a higher dose (500 nmol/kg) of GIP, suggesting that the observed loss of function was only partial (**Figure 2M-P**). These observations present across both sexes (**Figure S2** and **S3**).

### *Gipr^Halo/Halo^* pancreatic islet cells present impaired GIP-induced cAMP signalling

In view of the reduced anti-hyperglycaemic response to GIP in *Gipr^Halo/Halo^* mice, we analysed cAMP signalling dynamics in dispersed pancreatic islets from *Gipr^Wt/Wt^* and *Gipr^Halo/Halo^* mice transduced with the fluorescence-based cADDis cAMP biosensor and subsequently stimulated with a stepwise concentration gradient of human or mouse GIP (**Figure 3A-F**). In accordance with the *in vivo* glucose tolerance results, islets from *Gipr^Halo/Halo^* mice still responded to GIP, but there was a statistically significant reduction in EC_50_ compared to islets from *Gipr^Wt/Wt^* mice (mouse GIP EC_50_: *Gipr^Wt/Wt^* = 11.36 nM, *Gipr^Halo/Halo^* = 72.23 nM (p=0.01 by unpaired t-test of logEC_50_s); human GIP EC_50_: *Gipr^Wt/Wt^* = 42.43 nM, *Gipr^Halo/Halo^* = 146.78 nM (p=0.07 by unpaired t-test of logEC_50_s)).

**Figure 3:**
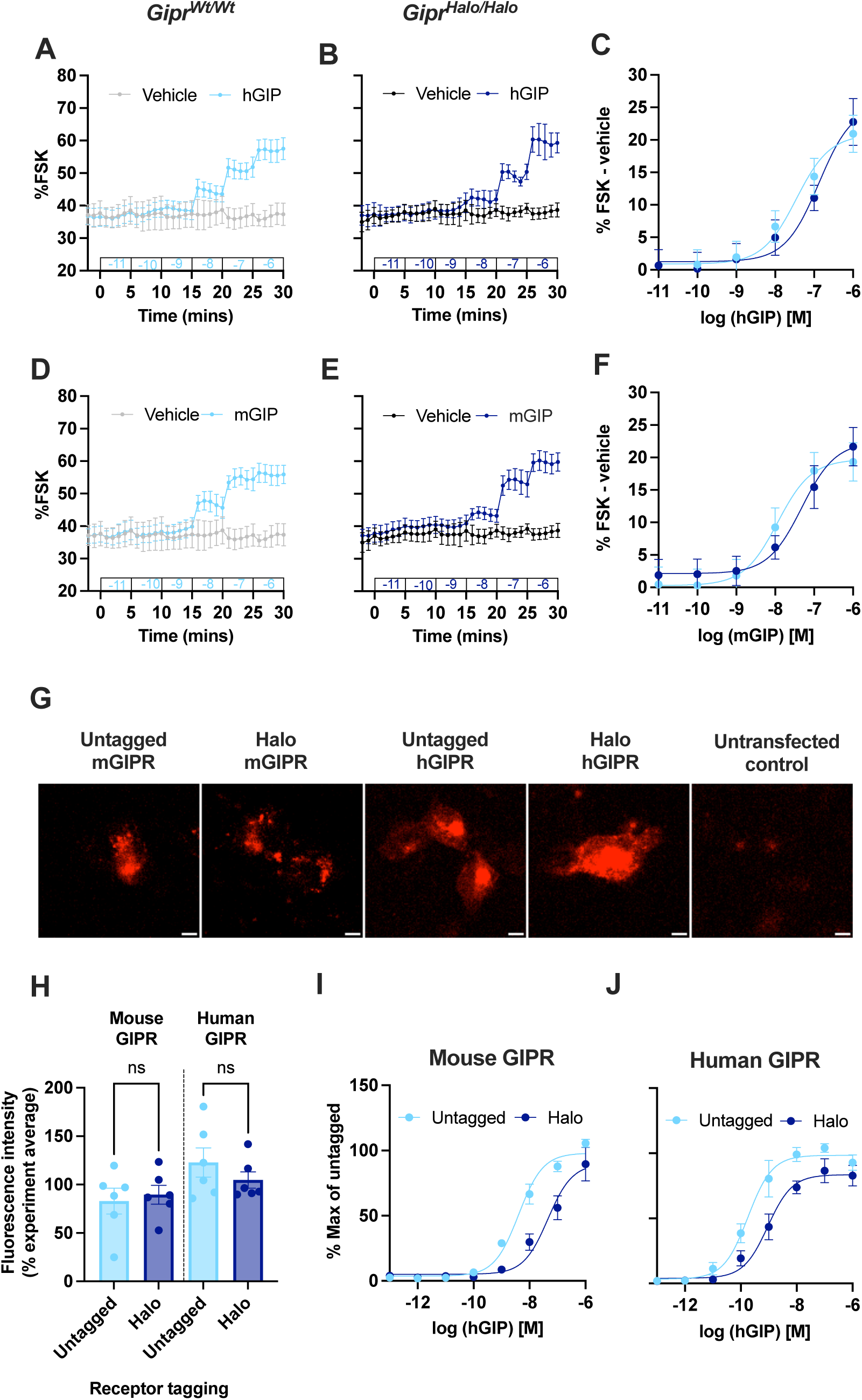
Islets from *Gipr^Halo/Halo^* mice generate cAMP in response to higher doses of human GIP and mouse GIP than *Gipr^Wt/Wt^* mouse islets. **A**-**F**) cAMP signal as a percentage of FSK/IBMX responses in cADDis-transduced dispersed islet cells from *Gipr^Wt/Wt^* mice (n=3-4) (**A**, **D**) or *Gipr^Halo/Halo^* mice (n=4) (**B**, **E**), in response to stepwise addition of human GIP (hGIP; **A**, **B**) and mouse GIP (mGIP; **D**, **E**). **C**, **H**) cAMP dose-response curves derived from AUC in each time interval in **A** and **B** (for **C**) and **D** and **E** (for **F**) minus vehicle control, with three parameter fits shown. **G**) Representative images of experiment displayed in **H** (scale bars = 20 µm). **H**) Fluorescence intensity of AD293 cells transiently transfected with mouse and human untagged and Halo-tagged GIPR following 1 hour of labelling with GIP-TMR (100 nM) (n=6), expressed as a percentage of the experimental average. Data was analysed using a one-way ANOVA. The Šídák test was used to correct for multiple comparisons. **I**, **J**) cAMP dose responses in AD293 cells transiently transfected with mouse (**I**) and human (**J**) untagged and Halo-tagged GIPR, stimulated for 30 minutes with increasing doses of hGIP (n=4-5). Three parameter fits are shown, with values normalised to the % max of the mouse or human untagged receptor. All values are presented as a mean ± SEM.

We then wondered if this impairment in cAMP signalling could be specific to endogenously expressed Halo-GIPR, which might be expected to reveal subtle differences in function not seen in heterologous systems. Therefore, we performed cAMP assays using AD293 cells transiently transfected with either untagged or Halo-tagged human or mouse GIPR, all driven by a CMV promoter. High content cell surface labelling with the fluorescent GIP analogue GIP-TMR (100 nM) suggested no impact of the Halo-tag on delivery of the human or mouse GIPR to the plasma membrane (**Figure 3G-H**) (13). However, in keeping with our *ex vivo* results from dispersed mouse islets, cells expressing the Halo-mGIPR displayed an impaired cAMP response to GIP compared to untagged mGIPR (untagged mGIPR EC_50_ = 5.97 nM, Halo-mGIPR = 55.52 nM, p=0.01 by paired t-test of logEC_50_s), a finding also observed at the human GIPR (untagged hGIPR EC_50_ = 0.36 nM, Halo-hGIPR = 1.63 nM, p=0.02 by paired t-test of logEC_50_s) (**Figure 3I-J**). Thus, these results suggest that in *in vitro* models, N-terminal Halo-tagging of the GIPR had not affected cell surface expression levels but had either affected ligand binding (at doses lower than 100 nM) or intracellular G protein signalling from the receptor.

### Visualising the Halo-tagged endogenous GIPR in pancreatic islets

As shown above, despite reduced potency, exogenously expressed Halo-GIPR was still expressed and trafficked to the cell surface in heterologous cells. We therefore tested whether our Halo-GIPR mouse model would still allow to study endogenous expression patterns of the GIPR by Halo-tag detection. We focussed on visualising the endogenous GIPR in pancreatic islets due to relatively high levels of *Gipr* expression in the pancreas compared to other tissues. While initial attempts to label islets using Halo-probes were unsuccessful, anti-Halo antibody staining revealed a significantly higher intensity of signal at the surface of islet cells in *Gipr^Halo/Wt^* islets compared to *Gipr^Wt/Wt^* islets (**Figure 4**). Although endogenous islet GIPR expression has previously been inferred via fluorescently tagged and radiolabelled agonists (4,19), including by our own studies (13), to our knowledge, this is the first evidence of direct detection of endogenous GIPR in pancreatic islets using antibody labelling.

**Figure 4:**
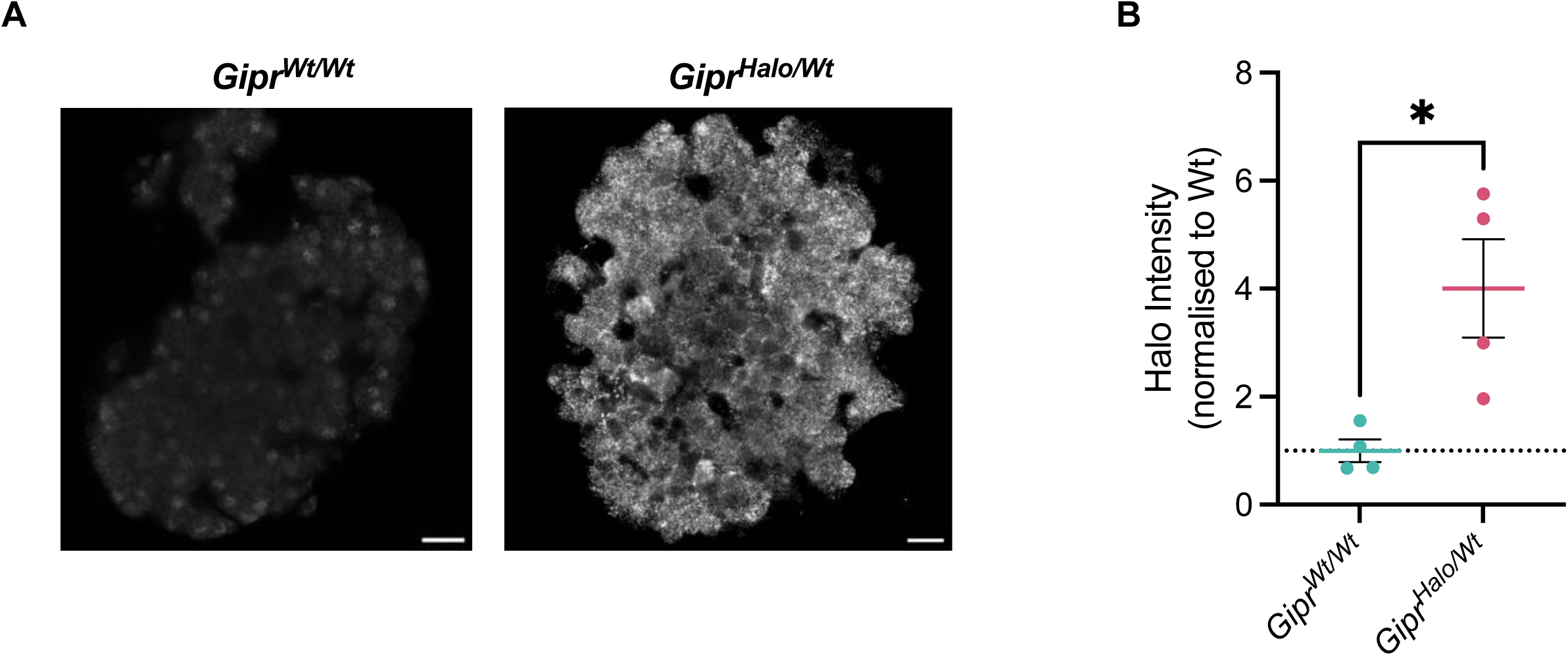
Endogenous GIPR visualised via anti-Halo antibody staining. **A**) Representative images of a pancreatic islet from *Gipr^Wt/Wt^* and *Gipr^Halo/Wt^* mice labelled with an anti-Halo antibody. Scale bar = 20 μm. **B**) Mean intensity of cell surface anti-Halo staining normalised to wildtype from 9-21 islets from *Gipr^Wt/Wt^* (n=4) *versus* littermate *Gipr^Halo/Wt^* (n=4) mice. Changes in fluorescence intensity were analysed using a student’s t-test. Values are presented as a mean ± SEM; *P<0.05.

Interestingly, we were unable to detect a signal in *Gipr^Halo/Halo^* islets (**Figure S4**). This suggests that endogenously expressing both *Halo-Gipr* alleles sufficiently alters receptor biosynthetic trafficking to prevent cell surface Halo-GIPR detection. This is a potential explanation of our *in vivo* and *ex vivo* observation of impaired Halo-GIPR function in response to GIP stimulation. Of note, this is at odds with our *in vitro* data which suggests that N-terminal Halo-tagging of the GIPR had not affected cell surface expression, but this was observed in an overexpressing system where partial trafficking defects might not be detectable.

## Discussion

In this project, we generated a *Gipr^Halo/Halo^* mouse model for the study of endogenous GIPR expression and trafficking. An *in vivo* characterisation of these mice showed that despite comparable body weights and oral glucose tolerance to *Gipr^Wt/Wt^* littermates, *Gipr^Halo/Halo^* mice only responded to high dose human GIP during IPGTTs, suggestive of impaired function of the N-terminally Halo-tagged receptor. Interestingly, despite physiological GIPR signalling being important for glucose handling in mice (20), glucose responses during an OGTT were normal in *Gipr^Halo/Halo^* mice. This could involve a potential upregulation of GLP-1R signalling to compensate for the germline impairment of GIPR function, as has previously been described in *Gipr* knockout mice (21).

*Ex vivo* assessment of islet GIPR function across littermates through cAMP assessment confirmed an impairment of Halo-tagged GIPR signalling. This was not unique to the endogenous GIPR, as the same result was observed when performing cAMP accumulation assays in AD293 cells transiently transfected with the wild-type *versus* the Halo-tagged GIPR. Interestingly, while surface GIPR expression was not impaired by Halo-tagging the receptor in an overexpressing system *in vitro*, anti-Halo staining in mouse islets showed detectable signal in *Gipr^Halo/Wt^* but not in *Gipr^Halo/Halo^* islets, suggesting that homozygous expression of endogenous Halo-tagged GIPR disrupted biosynthetic trafficking of the receptor to the cell surface. Thus, it appears that impaired Halo-GIPR signalling *in vivo* could be due to both reduced cell surface expression and impaired downstream signalling responses.

This study highlights that, despite their small size, adding enzyme self-labels to proteins risks altering protein function. This is not the first example of this phenomena; N-terminally Halo-tagging of the endogenous cannabinoid type 2 receptor (CB2) resulted in confirmational changes to the receptor which did not affect surface expression but prevented subsequent G protein recruitment (22). Endogenous SNAP-tagging of the other incretin receptor, GLP-1R, has been successfully achieved in mice, with SNAP-GLP-1R-expressing islets retaining signalling responses comparable to WT with 20 nM exendin-4 (12). This suggests the GLP-1R is more tolerant of the utilsed SNAP-tag than was GIPR of the Halo-tag in our study, although subtle signalling and trafficking alterations e.g. at non-saturating doses of GLP-1R agonists are still possible.

While Ast *et al.* were able to specifically label SNAP-GLP-1R-expressing islets with fluorescent SNAP-tag probes (12), we were unable to successfully label Halo-GIPR-expressing islets with similar Halo-probes. This could potentially be linked to the reduced endogenous GIPR surface expression compared to the GLP-1R in primary islets (13), which might have been potentially exacerbated by the addition of the Halo tag. While use of improved Halo probes was a possibility (16), we employed an antibody approach and were able to successfully visualise islet GIPR expression.

Therein, while our results call for caution when using *Gipr^Halo/Halo^* mice to study GIPR functional responses, our success in labelling endogenous GIPR in pancreatic islets suggests that this model could still be a valuable tool in evaluating GIPR expression in tissues where this is contentious, such as adipose tissue and the CNS. Whilst *Gipr*^EYFP^ mouse models have enabled cell sorting and subsequent transcriptomic analysis of hypothalamic and hindbrain cells with *Gipr* mRNA expression (3,4), our model allows for a similar analysis with the advantage enabling selection of cells in which GIPR protein expression is proven.

In conclusion, the addition of a Halo-tag to the endogenous GIPR resulted in an impairment of its signalling. This limits the use of the present mouse model to expression studies and is a cautionary tale to the use of enzyme-self labels in endogenous GPCR labelling.

## Acknowledgements

The Section of Endocrinology and Investigative Medicine at Imperial College London is funded by grants from the MRC, NIHR and is supported by the NIHR Biomedical Research Centre Funding Scheme and the NIHR/Imperial Clinical Research Facility. The views expressed are those of the author(s) and not necessarily those of the NHS, the NIHR or the Department of Health. The AT lab is funded by an MRC Project Grant (MR/X021467/1) and a Wellcome Trust Discovery Award (301619/Z/23/Z). ID is funded via a Medical Research Council Doctoral Training Partnership. TMMT is supported by the NIHR BRC and grants from MRC, DUK and NIHR. BJ acknowledges funding support from the Medical Research Council (MR/Y00132X/1 and MR/X021467/1), the Wellcome Trust (301619/Z/23/Z), Diabetes UK, the Eli Lilly and Company LRAP programme, and Metsera Inc. AT also acknowledges funding from Diabetes UK, the Society for Endocrinology and the Eli Lilly and Company LRAP programme.

The authors thank the ES Cell & Transgenic Facility of the Laboratory of Medical Sciences, Medical Research Council for the rederivation of *Gipr^Halo/Halo^* mice, Imperial College Facility for Imaging by Light Microscopy (FILM) for access to imaging facilities and technical support and Dr Aida Martinez-Sanchez and Prof Kevin G Murphy for providing animal project licence access.

