## Supplementary Information for "The physiological impact of an N-terminal Halo-tag on GIPR function in mice"

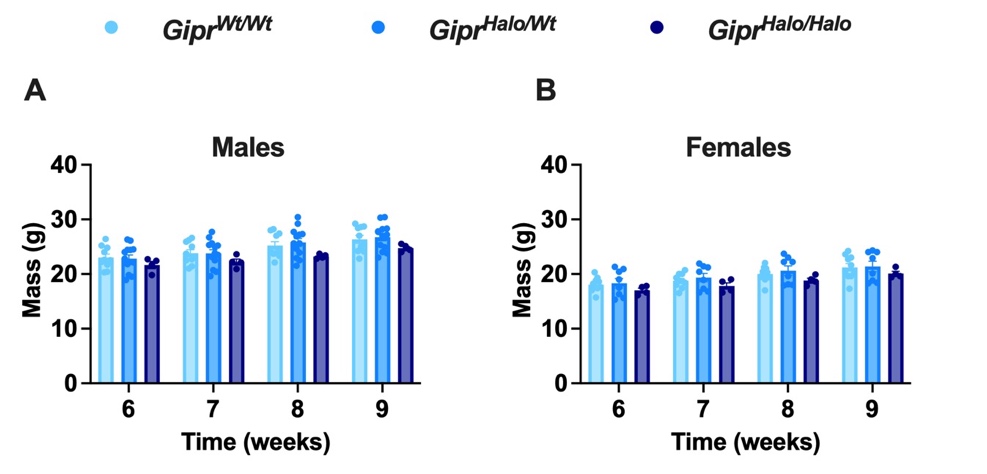

##### **Supplementary Figure 1: No difference in body weights between *Gipr^Wt/Wt^*, *Gipr^Halo/Wt^* and *Gipr^Halo/Halo^* mice at 6-9 weeks of age.**

**A**) Body mass (g) of *Gipr^Wt/Wt^* (male: n=10, female: n=10), *Gipr^Halo/Wt^* (male: n=13, female: n=8) and *Gipr^Halo/Halo^* (male: n=4, female: n=4), measured at 6-9 weeks of age. Males are displayed in **A** and females in **B.** Data was analysed using a two-way ANOVA with time and subgroup as co-variables. The Šídák test used to correct for multiple comparisons. Values are presented as a mean ± SEM.

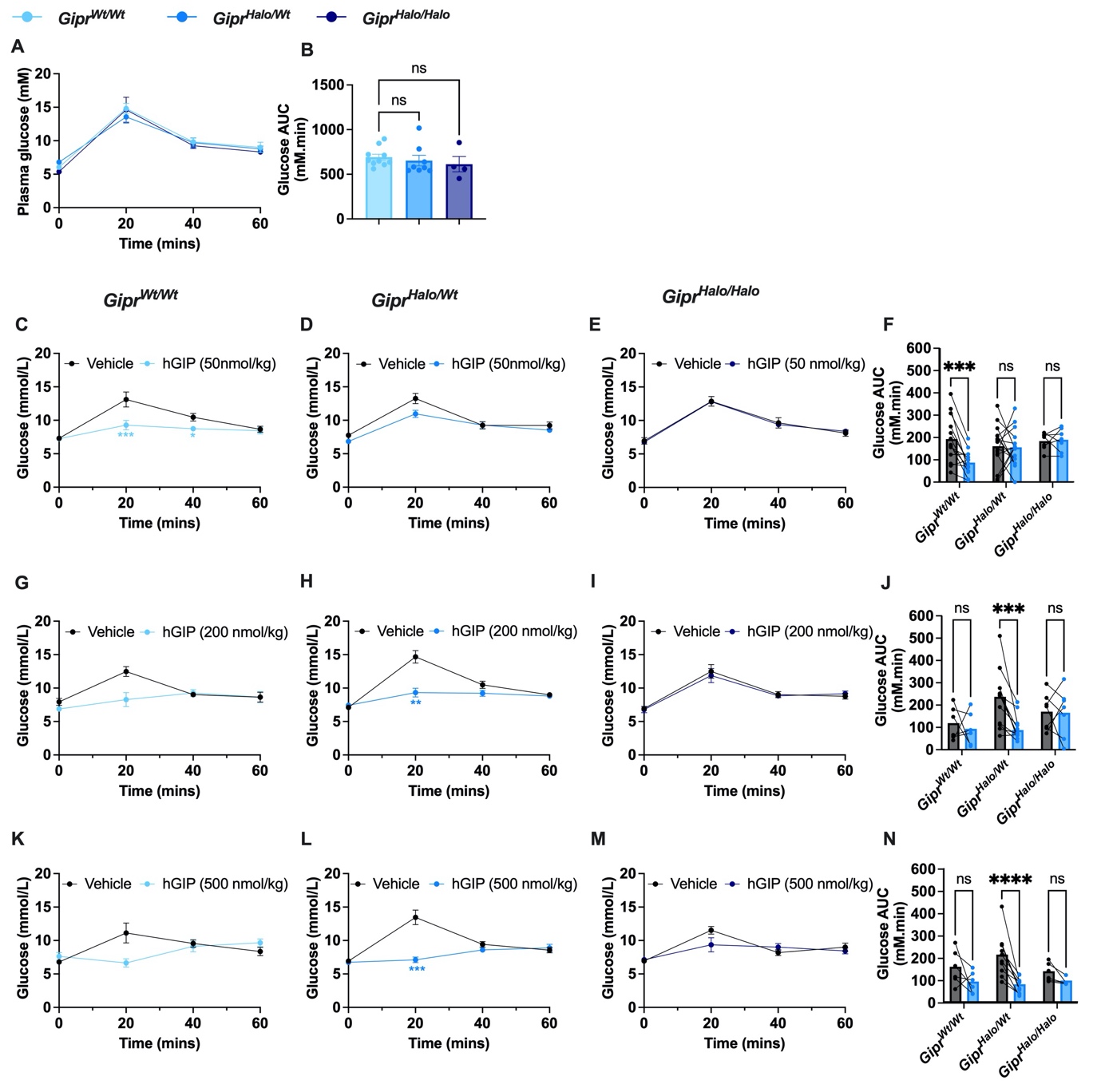

**Supplementary Figure 2: *Gipr^Halo/Halo^*** **female mice improve glucose tolerance in response to higher doses of human GIP than *Gipr^Wt/Wt^*** **female** **mice.**

**A**, **B**) OGTT conducted in *Gipr^Wt/Wt^* (n=10), *Gipr^Halo/Wt^* (n=8) and *Gipr^Halo/Halo^* (n=4) female mice. **C**-**N**) Crossover IPGTTs conducted in *Gipr^Wt/Wt^* female mice (**C**: n=13, **G**: n=6, **K**: n=6), *Gipr^Halo/Wt^* female mice (**D**: n=16, **H**: n=13, **L**: n=13) and *Gipr^Halo/Halo^* female mice (**E**: n=7, **I**: n=7, **M**: n=7), in response to human GIP (hGIP) 50 nmol/kg (**C**-**F**), hGIP 200 nmol/kg (**G-J**) and hGIP 500 nmol/kg (**K**-**N**). **A**, **C**-**E**, **G**-**I**, **K**-**M**) Plasma glucose time-course. **B**, **F**, **J**, **N**) Glucose AUC derived from corresponding glucose curves. Blood glucose at specific time-points have been analysed using a two-way ANOVA with time and subgroup as co-variables. The Šídák test was used to correct for multiple comparisons. Glucose AUC in **B** has been analysed with a one-way ANOVA. The Dunnett’s test was used to correct for multiple comparisons. Glucose AUCs in **F, J** and **N** have been analysed using a two-way ANOVA with genotype and subgroup as co-variables. The Šídák test was used to correct for multiple comparisons. All values are presented as a mean ± SEM. *P<0.05, **P<0.01, ***P< 0.001, ****P< 0.0001.

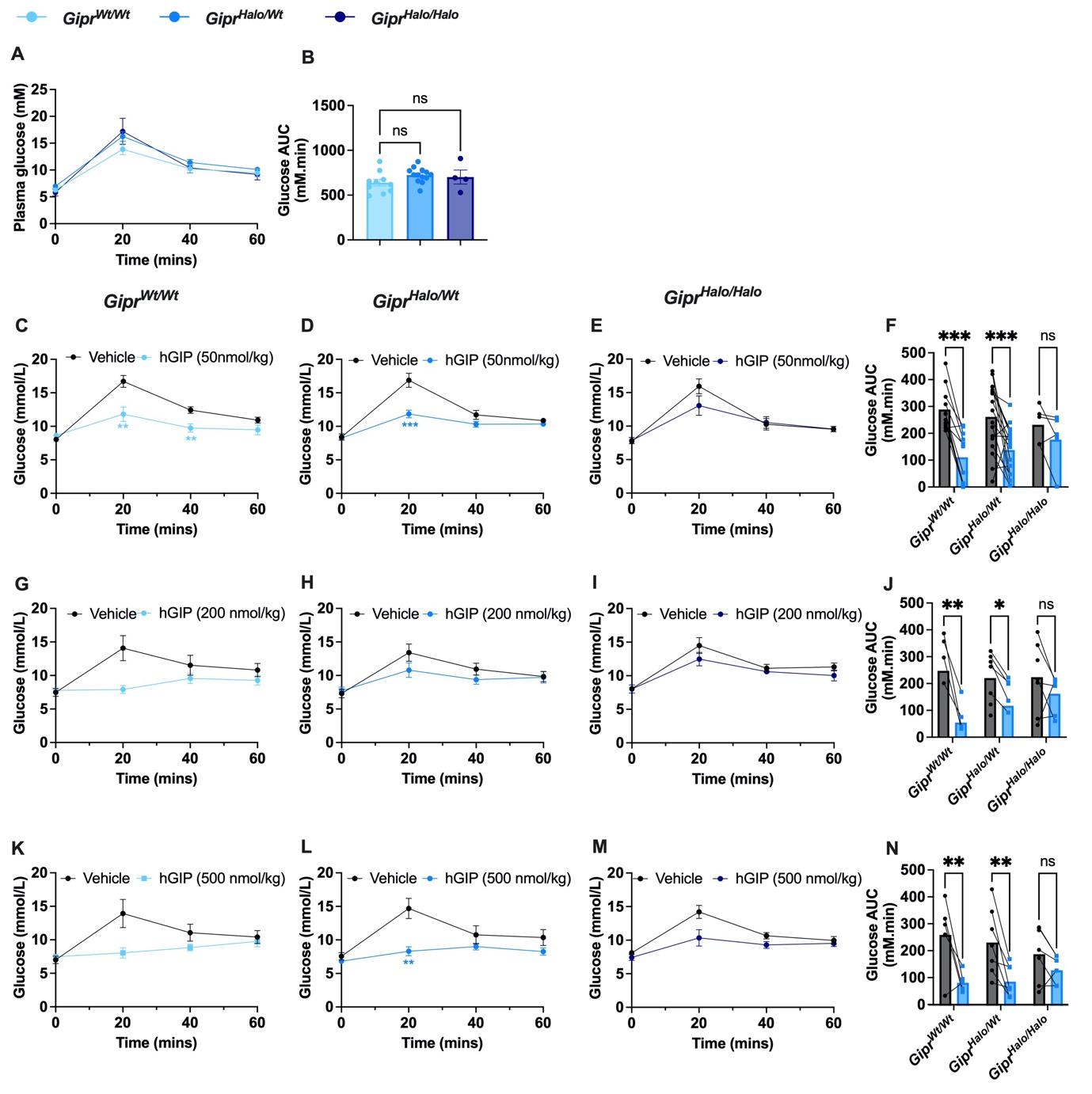

**Supplementary Figure 3: *Gipr^Halo/Halo^*** **male mice improve glucose tolerance in response to higher doses of human GIP than *Gipr^Wt/Wt^*** **male** **mice.**

**A**-**B**) OGTT conducted in *Gipr^Wt/Wt^* (n=10), *Gipr^Halo/Wt^* (n=13) and *Gipr^Halo/Halo^* (n=4) male mice. **C**-**N**) Crossover IPGTTs conducted in *Gipr^Wt/Wt^* male mice (**C**: n=11, **G**: n=5, **K**: n=5), *Gipr^Halo/Wt^* male mice (**D**: n=17, **H**: n=7, **L**: n=7) and *Gipr^Halo/Halo^* male mice (**E**: n=5, **I**: n=7, **M**: n=7), in response to human GIP (hGIP) 50 nmol/kg (**C**-**F**), hGIP 200 nmol/kg (**G**-**J**) and hGIP 500 nmol/kg (**K**-**N**). **A**, **C**-**E**, **G**-**I**, **K**-**M**) Plasma glucose time-course. **B**, **F**, **J**, **N**) Glucose AUC derived from corresponding glucose curves. Blood glucose at specific time-points have been analysed using a two-way ANOVA with time and subgroup as co-variables. The Šídák test was used to correct for multiple comparisons. Glucose AUC in **B** has been analysed with a one-way ANOVA. The Dunnett’s test was used to correct for multiple comparisons. Glucose AUCs in **F, J** and **N** have been analysed using a two-way ANOVA with genotype and subgroup as co-variables. The Šídák test was used to correct for multiple comparisons. All values are presented as a mean ± SEM. *P<0.05, **P<0.01, ***P< 0.001, ****P< 0.0001.

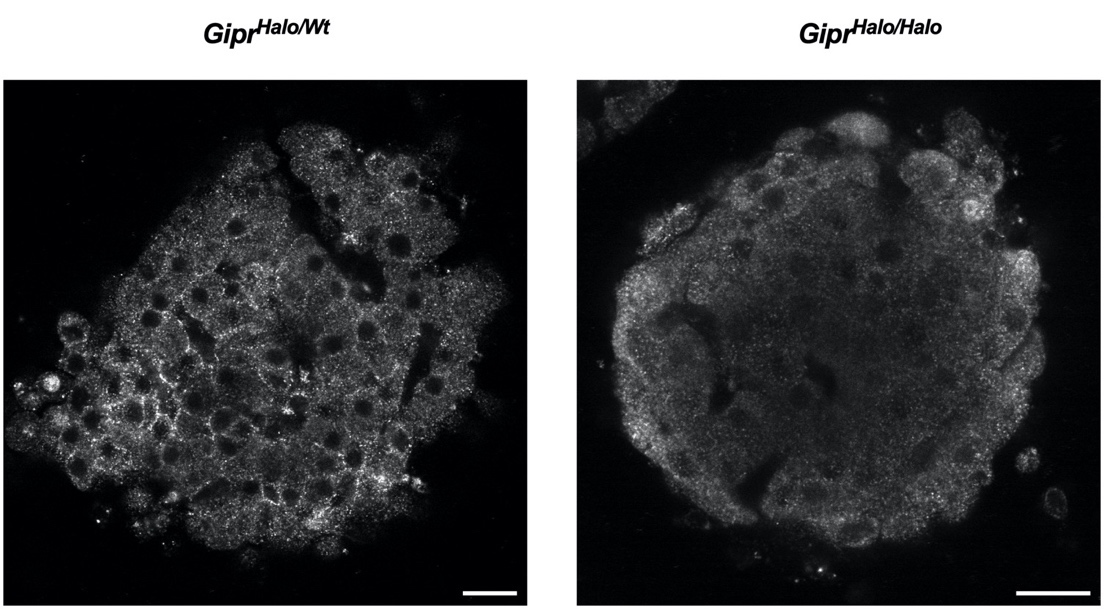

**Supplementary Figure 4: Anti-Halo staining in *Gipr^Halo/Halo^ versus Gipr^Halo/Wt^* mouse islets.** Representative images of a pancreatic islet from *Gipr^Halo/Wt^* compared to a *Gipr^Halo/Halo^* mouse islet labelled with an anti-Halo antibody. Scale bars = 20 μm.
